# Engineering pulsatile communication in bacterial consortia

**DOI:** 10.1101/111906

**Authors:** James M. Parkin, Victoria Hsiao, Richard M. Murray

## Abstract

Lux-type quorum sensing systems enable communication in bacteria with only two protein components: a signal synthase and an inducible transcription activator. The simplicity of these systems makes them a popular choice for engineering collaborative behaviors in synthetic bacterial consortia, such as photographic edge detection and synchronized oscillation. To add to this body of work, we propose a pulsatile communication circuit that enables dynamic patterning and long-distance communication analogous to action potentials traveling through nerve tissue. We employed a model-driven design paradigm involving incremental characterization of *in vivo* design candidates with increasing circuit complexity. Beginning with a simple inducible reporter system, we screened a small number of circuits varying in their promoter and ribosomal binding site (RBS) strengths. From this candidate pool, we selected a candidate to be the seed network for the subsequent round of more complex circuit variants, likewise variable in promoter and RBS strengths. The selection criteria at each level of complexity is tailored to optimize a different desirable performance characteristic. By this approach we individually optimized reporter signal-to-background ratio, pulsatile response to induction, and quiescent basal transcription, avoiding large library screens while ensuring robust performance of the composite circuit.

## 1 Introduction

Natural quorum sensing circuits implement positive feedback to establish bistable control over quorum sensing components. The feedback is mediated by the quorum sensing transcription factor, which activates transcription from quorum sensing promoters when bound to its cognate acyl-homoserine lactone (AHL) signal molecule. Activated transcription factor promotes synthase expression and, in turn, AHL accumulation. This feedback, along with rapid diffusion of AHL, ensures two stable states: one with only basal expression of quorum sensing-related expression and the other with high, saturated expression of quorum sensing components and maximal synthesis of AHL [11, 10]. In this paradigm, a quorum sensing circuit essentially contributes one bit of information: high or low density.

Augmenting the natural quorum sensing circuit with a negative feedforward arm forms an incoherent feedforward loop (IFFL), a well-studied circuit family capable of a variety of behaviors such as bistability, oscillation, and pulse generation [9, 3, 2]. Groundbreaking work in synthetic synchronized oscillators has established that, in accordance with the Turing model of reaction-diffusion systems, extending the spatial domain relative to the diffusion coefficient results in qualitatively different behaviors [13, 6]. Namely, a circuit that produces synchronized oscillations over small spatial domains will behave as a wave generator in larger arenas. In demonstrations involving extended arenas, it was shown that a single cluster spontaneously became the source of signaling activity for the entire community. This cluster emits, at regular intervals, signal packets traveling away from their source in all directions. Taking inspiration from these results, we aim to apply a similar IFFL gene circuit in a pulsatile communication circuit. Our goal is to tune the transcriptional balances such that there exists stable a low-expression state of the circuit, yet it will emit a single pulse in response to increased signal chemical concentration. This system would enable cells to initiate and transmit communications without permanently altering their internal transcriptional state or spontaneously generating signals. In this construction, a single quorum sensing system emulates a serial port, where a single channel may transmit more than one bit of information.

We hope that work on this system will open the door to more distributed logic and pattern formation applications in synthetic biology. Previous work has shown that dynamic transcriptional programs and heterogeneous populations may allow for more informative chemical event detectors, embedded control of cell growth, and photographic edge detection [5, 1, 12]. One of our goals for this research is to build an *E. coli* strain capable of detecting the temporal coincidence of spatially separated chemical events. As mentioned above, when used as a serial port, a signaling channel may communicate more than one bit of information. Specifically, a signal pulse generated in response to a single event carries information on the time of event in addition to the occurrence of the event itself. In a scenario with two such circuits, based on orthogonal quorum sensing components, the collision of pulses indicates that they were both emitted within a window of time that is dependent on the size of the consortia and the transmission velocity of the pulses.

## 2 Circuit design and modeling

The proposed circuit is an incoherent feedforward loop composed of a Lux-type quorum sensing system augmented with a negative feedforward arm. Lux-type quorum sensing systems consist a transcription factor, a signal molecule, and synthase enzyme, and a transcriptional repressor (Fig. 1). In this system, the quorum sensing transcription factor is expressed constitutively and, when bound to its cognate quorum sensing molecule, activates expression of the repressor and signal synthase. One regulatory loop is formed by the quorum sensing components, a positive-feedback loop wherein the quorum sensing signal molecule promotes its own production via transcriptional activation of its associated synthase protein. The transcriptional repressor acts on the synthase's promoter region, forming negative feedforward arm which is delayed relative to the positive feedback arm.

**Figure 1:**
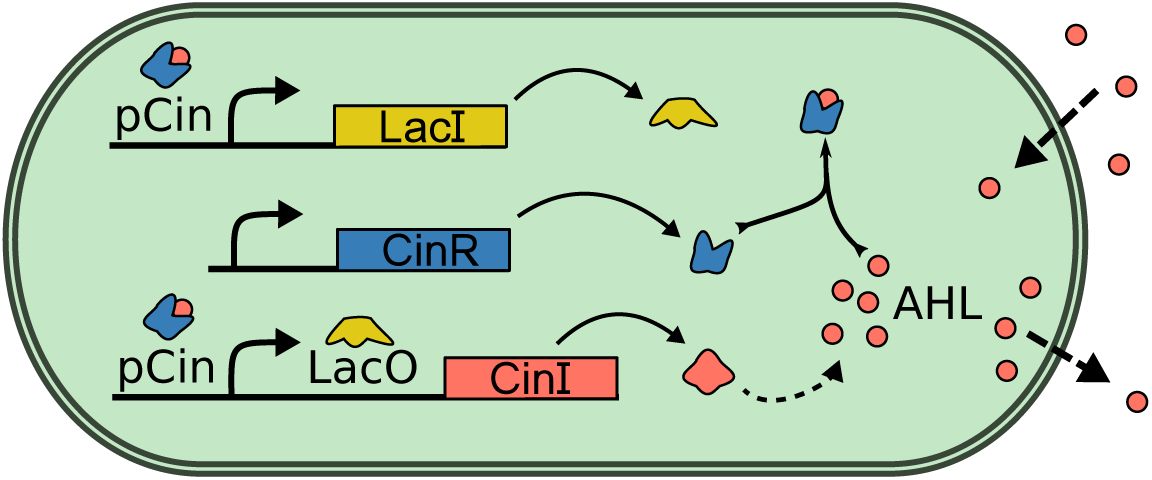
The schematic above depicts the protein, genetic, and chemical components of an example pulsatile communication circuit implemented with Cin quorum sensing components and LacI as the transcriptional repressor. The synthase protein, CinI, produces N-(3-hydroxy-7-*cis*-tetradecanoyl)-L-homoserine lactone (referred to as AHL in the context of the Cin system for convenience) [8]. CinR is expressed constitutively, AHL-bound CinR promotes expression from pCin, and LacI represses CinI transcription when bound to the LacO site. AHL freely diffuses through the cell wall.

Numerical simulations of the circuit show that it is capable of producing synchronized oscillations, bistability, or pulse transmission, among other behaviors (Fig. 2). Indeed, similar circuit architectures been applied successfully in synchronized oscillators [13, 3]. However, these behaviors are sensitive to variations in parameter values and accurately predicting transcriptional and enzymatic parameters from literature is haphazard [7]. To quickly search through design space, we build and characterize the circuit incrementally.

**Figure 2:**
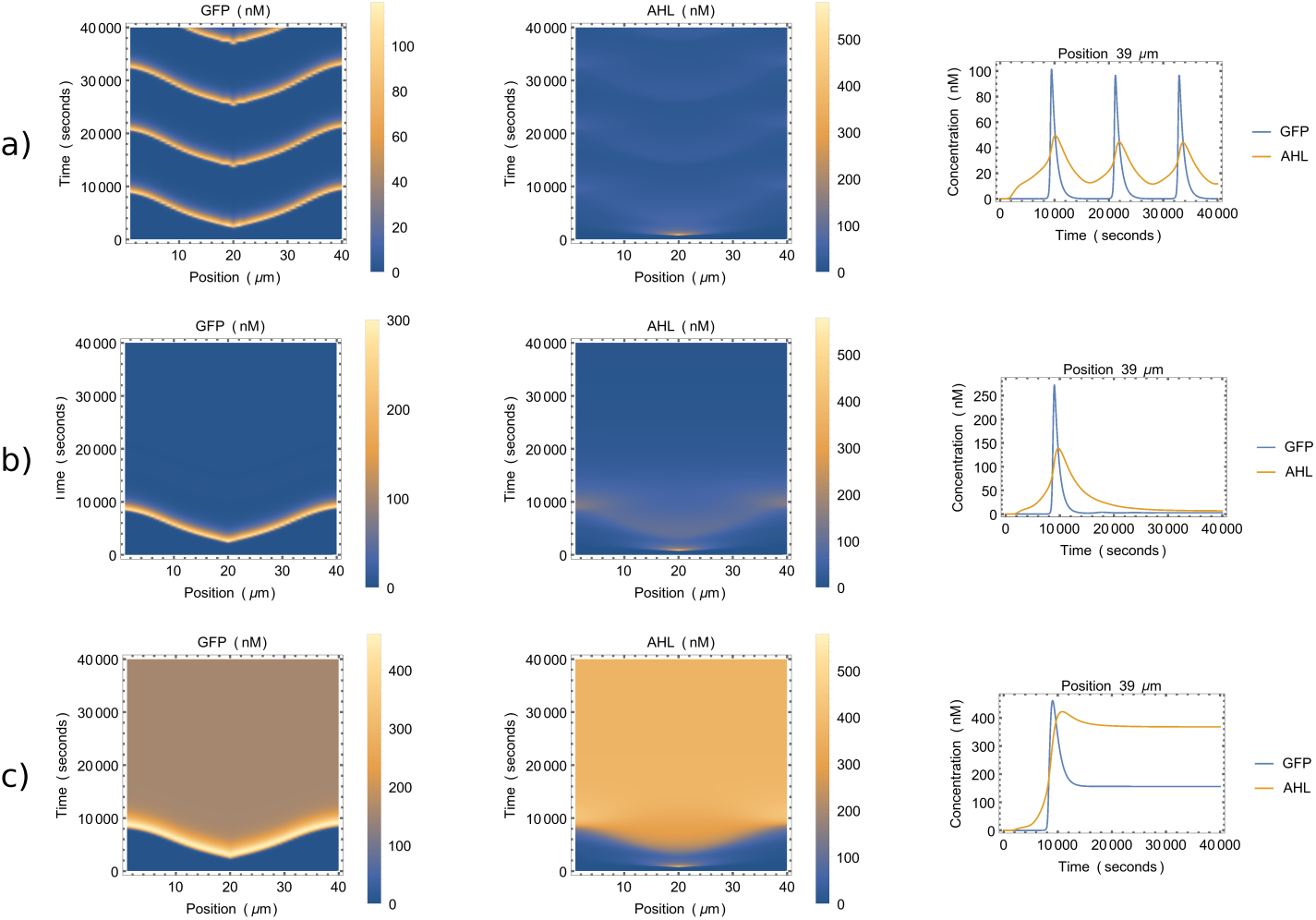
Simulations show behavior dependence on a single parameter *r_R_*, the maximum expression rate of the transcriptional repressor. In the model, GFP is under identical transcriptional control as the synthase protein and therefore indicates areas engaged in signaling. Left column plots show simulated GFP concentrations. Cells are homogeneously spread over one spatial dimension, indexed on the x-axis. Multiplication of the bacteria is not modeled, though biomolecule dilution due to division is modeled. Center column plots depict AHL concentration in the media and depict the experimenter-added AHL at time 800s and at location 20*μ*m. The line plots in the right column show concentration traces for both external AHL and intracellular GFP. **a)** *r_R_* = 25 yields oscillations in both concentrations. **b)** *r_R_* = 2.5 yields the desired behavior, a pulse of signaling activity that travels away from its source. **c)** *r_R_* = 0.25 produces bistable behavior, where the total cell population reaches a high-GFP state after induction with external AHL.

## 3 Model-guided design and experimental characterization

In our approach, we translate the reaction-diffusion model to a well-mixed solution model. Then, we perform liquid-culture *in vivo* experiments of many candidate gene circuits and fit the data to the well-mixed model. At each step, we define an optimality measure used to select an optimal candidate from the pool of candidate circuits to seed the following round of screening. In the following round, the seed circuit is augmented with variants of an additional component and the process is repeated.

Different optimality measures are required at each step. In the first round, we characterize variants of the constitutive quorum sensing transcription factor source and promoter variants of a fluorescent promoter and select for the highest inducible range of reporter expression. We add the repressor during the second round and select for the candidate whose maximal expression rate is largest with respect to its fully-repressed expression rate. In the final round, we add the synthase and select for the candidate that is quiescent when left alone and has a pulsatile response to externally provided AHL.

## 4 Experimental Results and Discussion

To date, we have experimentally characterized variations on the simplest circuit, an AHL-inducible fluorescent reporter. We compose these circuit candidates using two plasmids. One bears an inducible fluorescent protein and the other constitutively expresses a quorum sensing transcription factor. The reporter promoters are combinatorial, having activating operator sites for CinR binding and repressing operator sites for binding by either TetR or LacI. Experiments were performed using DH5a-Z1 *E. coli* as the host strain, which expresses both TetR and LacI from its genome. This way, cells began to express GFP at the beginning of the experiment, when aTc or IPTG were added.

The model predicts increasing intracellular GFP concentration at a constant rate during the first few hours of cell growth following induction by AHL along with IPTG or aTc. For each candidate, we determined its OD-normalized GFP expression rate in both 1*μ*M AHL and AHL-absent media [3]. The difference between these rates is the inducible range of expression rate, which is the optimality measure at this stage. The results (Fig.3) show that **J23106 B0032 CinR** yields high inducible range when paired with either **pCinTetO B0032 GFP** or **pCinLacO_m B0034 GFP** reporters (Table 1). Two reporters genotypes were selected so that we may test both TetR and LacI as repressors in the second round of screening.

**Figure 3:**
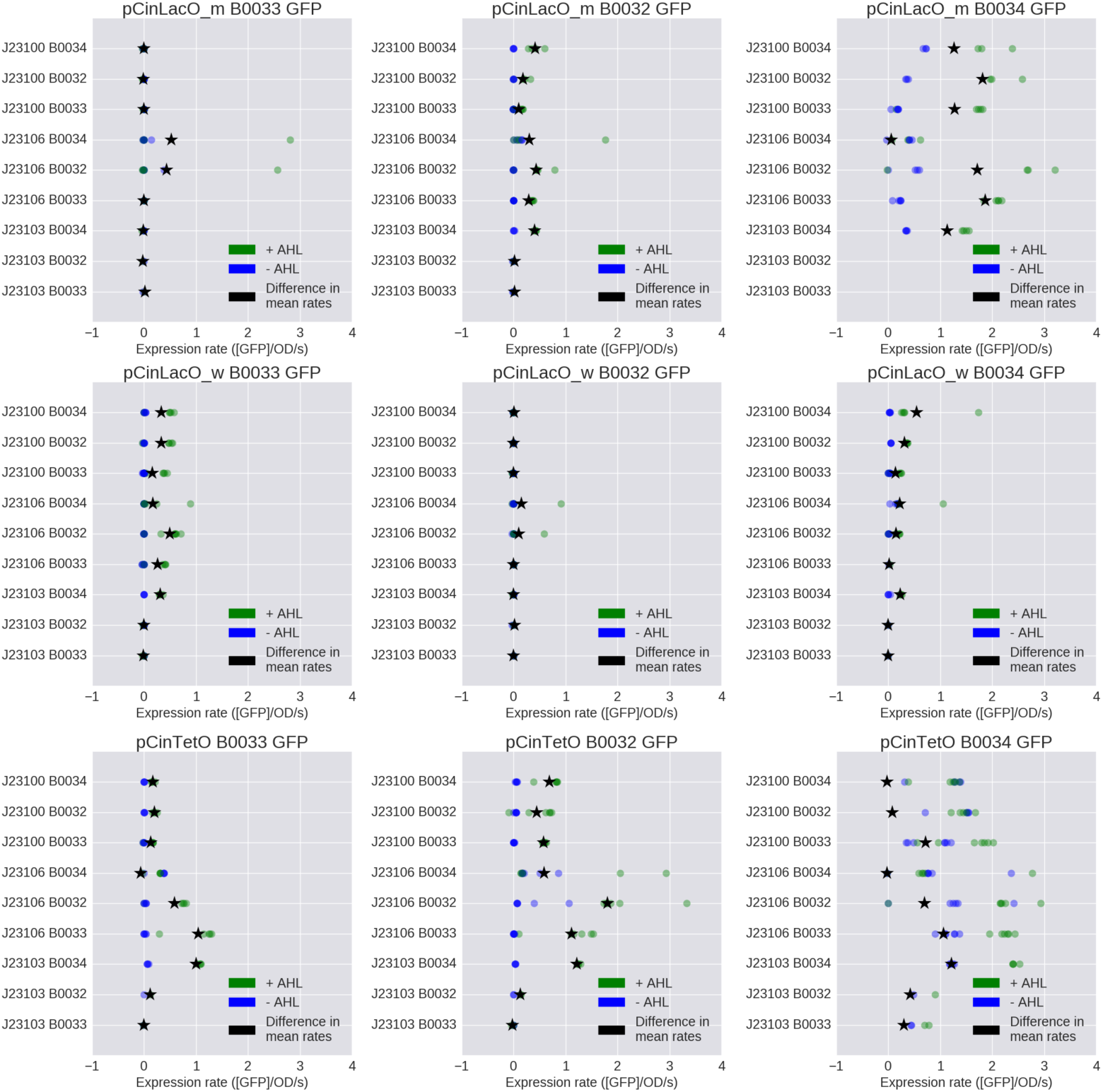
The OD-normalized GFP fluorescence from each experimental trial during the first round of screening was fit to a line. The slope of a fit line estimates the expression rate of GFP in that sample. In the plots above, the y-axis is labeled with the promoter and ribosomal binding site components that constitute the 5'UTR to the CinR coding region (See Table 1 for component descriptions). The titles specify the genotype of the fluorescent reporter plasmid. In each plot, the expression rates from no-AHL (blue dots) and high-AHL (green dots) trials are plotted along with the mean difference between these groups (black stars). Each genotype's difference of mean rates is its optimality value, and it appears that **J23106 B0032 CinR** pairs well with both **pCinTetO B0032 GFP** and **pCinLacO_m B0034 GFP** reporters.

**Table 1:** The genetic components used in this work were from the CIDAR MoClo Library kit. The table defines the promoter and ribosomal binding site (RBS) relative strengths as specified in the kit [4].

| Promoter | Strength | RBS | Strength |
| --- | --- | --- | --- |
| J23103 | Weak | B0033 | Weak |
| J23106 | Medium | B0032 | Medium |
| J23100 | Strong | B0034 | Strong |

For the second round of screening, a transcriptional repressor is added to the circuit architecture and variants are screened for pulsatile response to induction. In our first attempt, the repressor gene was inserted immediately downstream of the constitutively-expressed CinR. Preliminary screening data from shows almost no reporter repression resulting from the addition of a transcriptional repressor. We know that the promoters are repressible, having observed that repressors expressed from the DH5a-Z1 genome effectively suppress reporter transcription. In future work, we will investigate whether compositional context negatively impacts repressor expression, and can be ameliorated by reorganizing the gene allocation.

In addition, we will investigate how employing plasmids with different origins of replication can improve the performance and repeatability of the circuit. During the first stage of screening, plasmids pairs had different resistance genes but the same origin of replication, pMB1. Transforming two plasmids with related replication origins may result in heterogeneous plasmid copy number. Future work will include testing plasmid pairs with compatible pairs of replication origins.

## 5 Acknowledgments

James M. Parkin is supported by the Institute for Collaborative Biotechnologies through grant W911NF-09-0001 from the U.S. Army Research Office. The content of the information does not necessarily reflect the position or the policy of the Government, and no official endorsement should be inferred. Plasmid vectors and non-coding regions were provided as a generous gift of Douglas Densmore at the Cross-disciplinary Integration of Design Automation Research lab (Addgene Kit # 1000000059). Quorum sensing promoters and coding sequences were provided as a generous gift from Matthew Bennet (Addgene Plasmid # 65954, 65952).

